# Simplifying complexity: the reliability of lidar-derived forest structural complexity metrics for conservation management

**DOI:** 10.64898/2026.10.23.760726

**Authors:** Eleanor Clare Kent, William Rupert Moore Flynn, Phillippa R. Gullet, David Coomes, Emily Rebecca Lines

## Abstract

1. Forest structural complexity is regarded as a driver of biodiversity. The re-structuring of forests to increase structural complexity is now a common objective of forest restoration and conservation projects. Lidar provides an efficient means of monitoring forest structure; however, the reliability of lidar-derived structural complexity metrics remains uncertain. Most metrics require a choice of resolution (such as pixel or voxel size), and whilst metric outcomes are sensitive to this choice, an understanding of how this impacts subsequent ecological interpretations of forest restoration monitoring is missing.
2. We assess how metric resolution choices impact FSC estimates using terrestrial laser scanning data from 32 single-species forest plots in Cairngorms National Park, Scotland. These plots represent four contrasting structural categories: ancient woodland (AW), old plantation (OP), old plantation with re-structuring management (OPM) and natural restoration (NR); with a hypothesised order of complexity: AW, OPM, OP, NR (from highest complexity to lowest). We computed 21 FSC metrics and calculated how each metric ranked our forest structural categories across a resolution gradient (0.02-1.5m). We evaluated the stability of metric rankings across resolution values, their ability to discriminate forest categories, and the consistency of ecological interpretation of forest management targets.
3. Nine metrics produced unstable rankings, indicating that ecological conclusions were sensitive to resolution selection. Stable metrics discriminated well between categories with obvious structural differences but failed to discriminate between categories with subtle differences, particularly OP and OPM.
4. AW did not consistently exhibit the highest structural complexity, instead occupying an intermediate position characterised by a combination of canopy occupancy, openness and height inequality, not captured by any single metric.
5. We conclude that methodological choices can influence FSC estimates and ecological conclusions drawn from them.
6. *Synthesis and applications:* Reliable assessment of forest restoration depends not only on the choice of metric but also on how that metric is parameterised. We recommend transparent reporting and justification of resolution parameter selection together with the use of stable metrics that capture structural characteristics relevant to management objectives being monitored, to improve reliability and comparability of lidar-based forest monitoring.

## Introduction

Forest structural complexity (FSC) describes the diversity, abundance and spatial distribution of forest features in space (McElhinny et al., 2005). Heterogeneous environments provide a greater diversity and abundance of ecological niches that benefit organisms of different sizes, ages, and functional roles (Coverdale & Davies, 2023; LaRue et al., 2019). The increased vertical and horizontal variation in vegetation, tree crown dimensions, canopy gaps and deadwood provide diverse and environmentally variable habitats for foraging, breeding and shelter in relatively close proximity (McElhinny et al., 2005; Regnery et al., 2013; Zellweger et al., 2013). Consequently, structurally complex forests are often associated with higher levels of biodiversity, with positive associations described for birds (Carrasco et al., 2019), bats (Blakey et al., 2017), arthropods (Wildermuth et al., 2024), plants (Anderson et al., 2021) and fungi (Lang et al., 2023). Together, these relationships indicate that structural and biological diversity are closely interconnected, with structural variation influencing ecological processes and ecosystem functioning (Gao et al., 2014; LaRue et al., 2023; Martin et al., 2023; Pommerening, 2002).

Old growth forests in particular are often characterised by a high FSC resulting from long periods of natural disturbance, gap dynamics and stand development (Franklin & Van Pelt, 2004; Kuuluvainen, 2009), making them valuable reference system for forest restoration and conservation projects. Therefore, enhancing FSC in otherwise homogenous stands, such as old plantations, has become an increasingly important objective of forest restoration and management (Larsen et al., 2022). These interventions are typically designed to restore the natural conditions and ecological processes characteristic of complex old forests by emulating natural disturbance dynamics (Franklin & Van Pelt, 2004; Kuuluvainen, 2009). Common approaches include the retention of standing and fallen deadwood (The Enhancing Structural Complexity Project, Germany: Asbeck *et al*., 2023), the rewilding of large herbivores to influence regeneration dynamics and contribute to heterogeneous mosaics of open habitat, scrub and woodland (Kirby, 2004; Vera et al., 2006; Wilder Blean Project, Kent Wildlife Trust, n.d.) and the creation of canopy gaps, retention of veteran trees, and encouragement of multilayered canopy development (Mönkkönen et al., 2026; Zeller et al., 2023; Cairngorms Connect, n.d.). As the implementation of such conservation initiatives continues to expand, there is an increasing need for robust methods capable of monitoring the progress of FSC formation for biodiversity conservation forest restoration purposes.

Recent advances in high resolution remote sensing offer the opportunity to quantify a wide range of structural characteristics of forests directly and non-destructively (Calders et al., 2020; Disney, 2019; Lines et al., 2022). Light detection and ranging (lidar), an active remote sensing technology, generates high resolution, three-dimensional representations of a forest ecosystem. Lidar can be deployed from terrestrial and mobile scanning platforms on the ground (TLS and MLS), airborne platforms (airplane and drone), and spaceborne platforms such as NASA’s GEDI mission, enabling structural assessments across individual tree, landscape, and global scales (Calders et al., 2020; Disney, 2019; Lines et al., 2022). Among these, TLS provides the highest resolution point cloud data, capturing spatially referenced structural information from herb layer to canopy, often at millimetre accuracy (Disney, 2019). This level of detail enables the measurement of complex structural elements that are difficult or impossible to obtain with traditional field-based methods, including crown volume, leaf spatial arrangement, fractal space-filling properties, and foliage clumping (Béland & Baldocchi, 2020; Calders et al., 2020; Seidel, 2018). Consequently, lidar has become a popular tool for quantifying FSC and its relationship with biodiversity, including small mammals (Deere et al., 2020; Schooler & Zald, 2019), birds (Melin et al., 2018), and arthropods (Wildermuth et al., 2024). As lidar sensors continue to become smaller, more portable and cost-effective, they are being adopted more widely by researchers, foresters and conservation practitioners, offering opportunities for larger, landscape-scale understanding of FSC as well as national forest inventory surveys (Holvoet et al., 2025).

Despite widespread use, the quantification and interpretation of lidar-derived FSC estimates lacks standardisation. Numerous metrics have been developed in recent decades from a variety of mathematical and conceptual frameworks. Some intend to provide a holistic measure of FSC, such as box dimension, a fractal analysis metric that uses self-similarity assessment of forest features across multiple scales (Seidel, 2018), however, FSC is usually described using a collective suite of singular metrics that capture different structural characteristics (Table 1). Conceptually, these metrics can be grouped by dimension, as describing: vertical, horizontal or three-dimensional complexity, or into descriptive methodological groupings such as those proposed by Feigl et al., 2025 or Atkins et al., 2023a. The derivation of these FSC metrics often requires user-defined resolution parameters, such as pixel resolution a canopy height models (CHMs) or choice of voxel size or number of vegetation layers. Resolutions of these parameters can be selected pragmatically, either according to point cloud density (Li *et al*., 2017), with airborne or drone studies often use larger voxel or pixel sizes than TLS due to inherently lower point densities (e.g. 1m; Ross *et al*., 2024, 5m; Zhang *et al*., 2017, Table S2 in Supporting Information), or because resolution influences other sources of measurement uncertainty, such as occlusion (Ehbrecht et al., 2016). The proliferation of methods to quantify FSC has greatly improved the ability to assess the outcomes of conservation restructuring initiatives, but emerging in parallel is an acknowledgment of the lack of a unified, robust and scale-aware methodological standard across the literature (Loke and Chisholm, 2022; Cimdins *et al*., 2025), making standardisation and comparison of FSC a challenge.

**Table 1:** 21 metric descriptions split by resolution parameter and group as described by Feigl *et al*., 2025. Metrics are numbered 1 – 21 with the codes used in analysis for each metric in brackets proceeding the initial name.

| Group | Metric Name | Method description | Resolution parameter |
| --- | --- | --- | --- |
| Density | (1) Gap Fraction (3D) | Empty voxels / total voxels within the bounding box for each point cloud. | Voxel |
|  | (2) Vertical + horizontal space filling | The fraction of filled vs unfilled voxels at each height threshold summed vertically (SFz) with inequality statistics (3)(4) *Gini coefficient* (GINlz, GINlxy), and horizontally (5)(6) *Coefficient of Variation* (CVz, CVxy) and (Stiers *et al.*, 2020). Thresholds determined as the height/width of each voxel size. | Voxel |
|  | (7) CV of Leaf Area Density (LAD) | The coefficient of variation of total leaf area per unit volume at each height layer, derived from the vertical attenuation of LiDAR returns using the Beer-Lambert law, which corrects for signal occlusion (Bouvier *et al.*, 2015). Computed in the *leafR* package in R (R Core Team, 2025). | Voxel |
|  | (8) Effective Number of Layers (ENL) | Quantifies how many vertically distinct layers a forest effectively has, accounting for how vegetation is distributed among those layers (Ehbrecht *et al.*, 2016), using inverse Simpson index. Function from the *tre3d* R package (Kunz, 2019) | Voxel |
|  | (9) Foliage Height Diversity (FHD) | The Shannon Entropy of vegetation distribution across vertical strata (MacArthur & MacArthur, 1961). Computed from leaf area density (LAD) profiles using the *leafR* (Almeida *et al.*, 2021). | Height bin, voxel |
|  | (10) Vertical Complexity Index (VCI) | A measure of evenness, how evenly vegetation is distributed across layers within a forest plot. Calculated in R using the *lidR* package (Roussel *et al.*, 2020) Unlike the FHD, VCI is normalized by the maximum possible entropy and therefore reflects relative rather than absolute vertical complexity (Van Ewijk *et al.*, 2011). | Height bin |
| Fractal | (11) Box Dimension (BoxD) | The linear relationship between the logged total number of boxes needed to contain the point cloud and the log of box size (Seidel, 2018). Calculated using the *rTwig* package in R (Morales & MacFarlane, 2025). | N/A |
| Height | (12) Height | (12) mean and (13) max height | CHM |
| Rugosity | (14) Height complexity | (14) standard deviation, (15) coefficient of variation and (16) Gini coefficient of height, calculated directly from the CHM. | CHM |
|  | (17) Rumple index | The roughness of a surface as the ratio of that surface area compared with a flat plane. i.e. normalised ground. Computed using the *rumple_index* function in *lidR* (Roussel *et al.*, 2020) directly from a CHM. | CHM |
|  | (18) Rugosity | Mean standard deviation of height in a 3x3 moving window of a chm | CHM |
| Open-ness | (19) Gap fraction<br>(20) Canopy cover,<br>(21) CV of gap area | Proportion plot area that contains gaps or canopy cover above a relative threshold of 33% of z (shrub layer). Using package *ForestGapR* (Silva *et al.*, 2019) | CHM |

Evidence indicates that lidar-derived metrics are sensitive to user-defined resolution choices of metric parameters (Atkins et al., 2023b). Studies that explore metric sensitivity have used methods that focus on agreement with ground-truth measurements or comparisons between alternative platforms, often reporting inconsistent and context-dependent results (Atkins et al., 2023b; Chen et al., 2022; Cifuentes et al., 2014; Cimdins et al., 2025; Pimont et al., 2018). These studies provide valuable methodological contributions but do not necessarily address the consequences of drawing ecological conclusions from resolution-sensitive metrics. Consequently, when differences in FSC are detected, it is unclear whether differences are due to ecologically meaningful variation, or a product of methodological decisions (Pimont et al., 2018).

Here, we evaluate metric sensitivity to user-defined resolution choices, the impact of these choices on ecological interpretation and provide insights for broad and reproducible monitoring of forest restructuring management. We test the ability of multi-scan, stand-based FSC metrics, commonly found in the biodiversity literature, to reliably monitor structure across structurally diverse forest plots within the Cairngorms National Park, Scotland. We selected plots within active restoration landscape allowing us the opportunity to assess management interventions designed to restore plantation woodland to a more complex, resilient forest and chose plots of predominantly single species (*Pinus sylvestris*), to limit the confounding influence of tree species diversity on structure. This site is part We calculate 21 TLS-derived FSC metrics for 32 *Pinus sylvestris* plots, representing four contrasting categories of forest structure: Ancient Woodland (AW), old plantation (OP), old plantation with re-structuring management (OPM) and natural regeneration (NR). We evaluate FSC metrics according to three core criteria: 1) their *stability*, assessed as the consistency of FSC ranking order of categories across a gradient of resolution change; 2) their *discernability,* assessed as the ability of metrics to discriminate between forest categories, and 3) their ecological interpretation in relation to forest re-structuring management. We hypothesise that the plots within each forest structure category will exhibit differing structural complexity results, with a hypothesised order of complexity, from highest to lowest: AW, OPM, OP, NR.

## Methods

### Study site

Our study site is in the Cairngorms Connect Partnership area, which lies within the Cairngorms National Park (NP), in the Highlands of Scotland (Figure 1). The main soil type is podzol and conifers (predominantly the native *Pinus sylvestris*) have been the dominant species for at least 7000 years (Gauld, 1982; R. Summers, 2018). Forest cover declined substantially from around 6000 BP as forests were cleared for agriculture, grazing and timber extraction, reaching a historic low by the early twentieth century (5%; Peterken, 1983; Pratt, 2007). Today, the forests of the Cairngorms comprise ancient pinewood remnants, naturally regenerating woodland and old plantations, that are increasingly managed for conservation (Mason et al., 2007; Summers, 2018; Summers et al., 1997). Forest covers just 14.4% of the Cairngorms NP with approximately 56.0% comprising of old plantation (Amphlett, 2022).

**Fig. 1.**
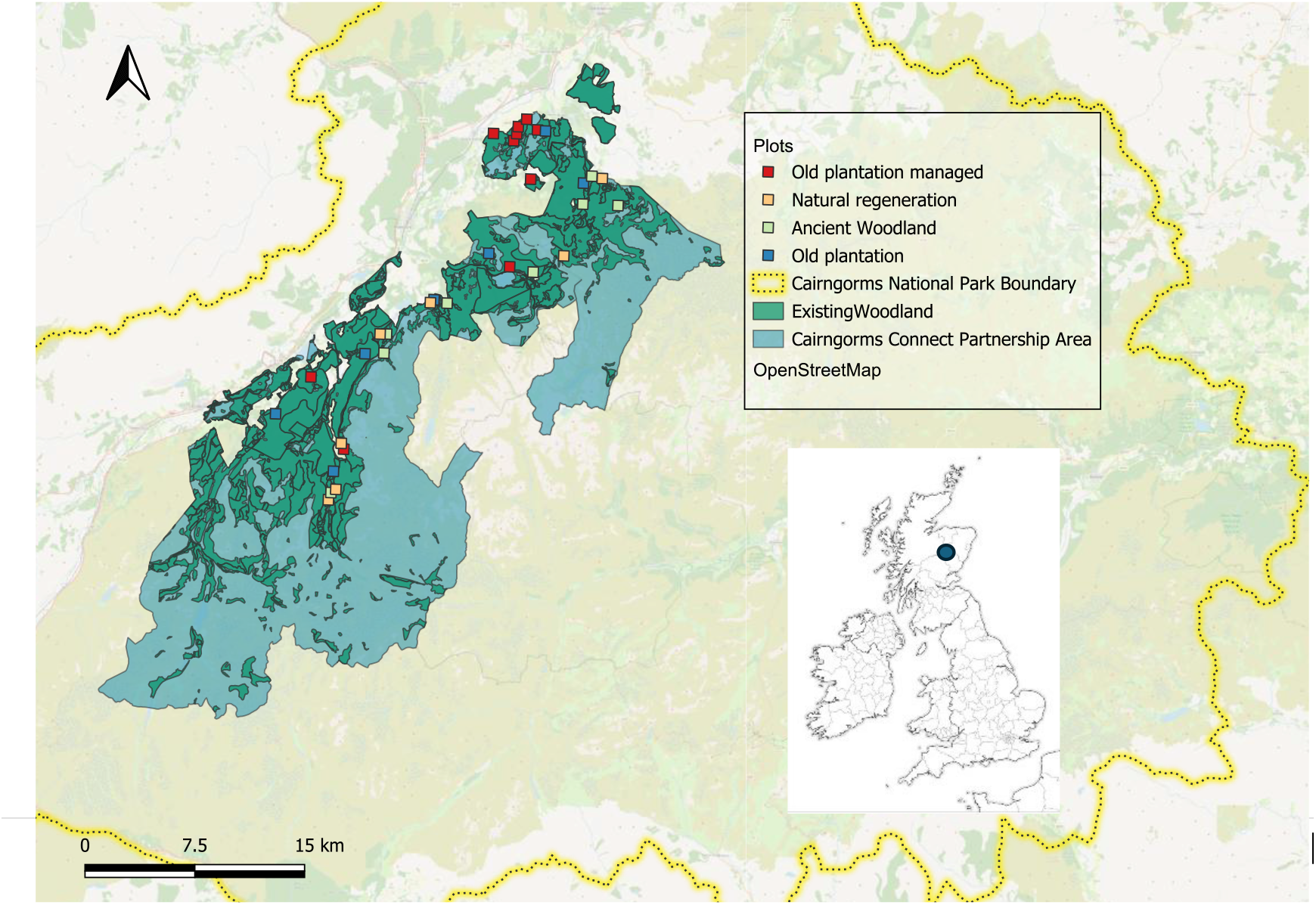
The Cairngorms Connect area (light green) and current woodland extent (blue), within the Cairngorms National Park boundary (yellow). Coloured squares show our 32

The Cairngorms Connect Partnership area is the largest contiguous restoration project in the UK, covering approximately 600 km² of the Cairngorms NP. Founded in 2014, Cairngorms Connect is a partnership between five organisations: Forestry and Land Scotland, NatureScot, RSPB Scotland, WildLand and the Cairngorms National Park Authority. The partnership’s goal is to restore native habitats, species and natural processes over the next 200 years, to deliver benefits for wildlife and people (Gullett et al., 2023).

### Forest Structural Categories

We identified four distinct forest structural types in the Cairngorms Connect landscape: Ancient Woodland (*AW*), old plantation (*OP*), old plantation managed for conservation (*OPM*) and natural regeneration (*NR*) (Figure 2). We selected plot locations after discussion with the Cairngorms Connect Partnership land managers and staff about land use history and recent/upcoming management plans for the area. All plots were between 215 – 418m elevation.

**Fig. 2:**
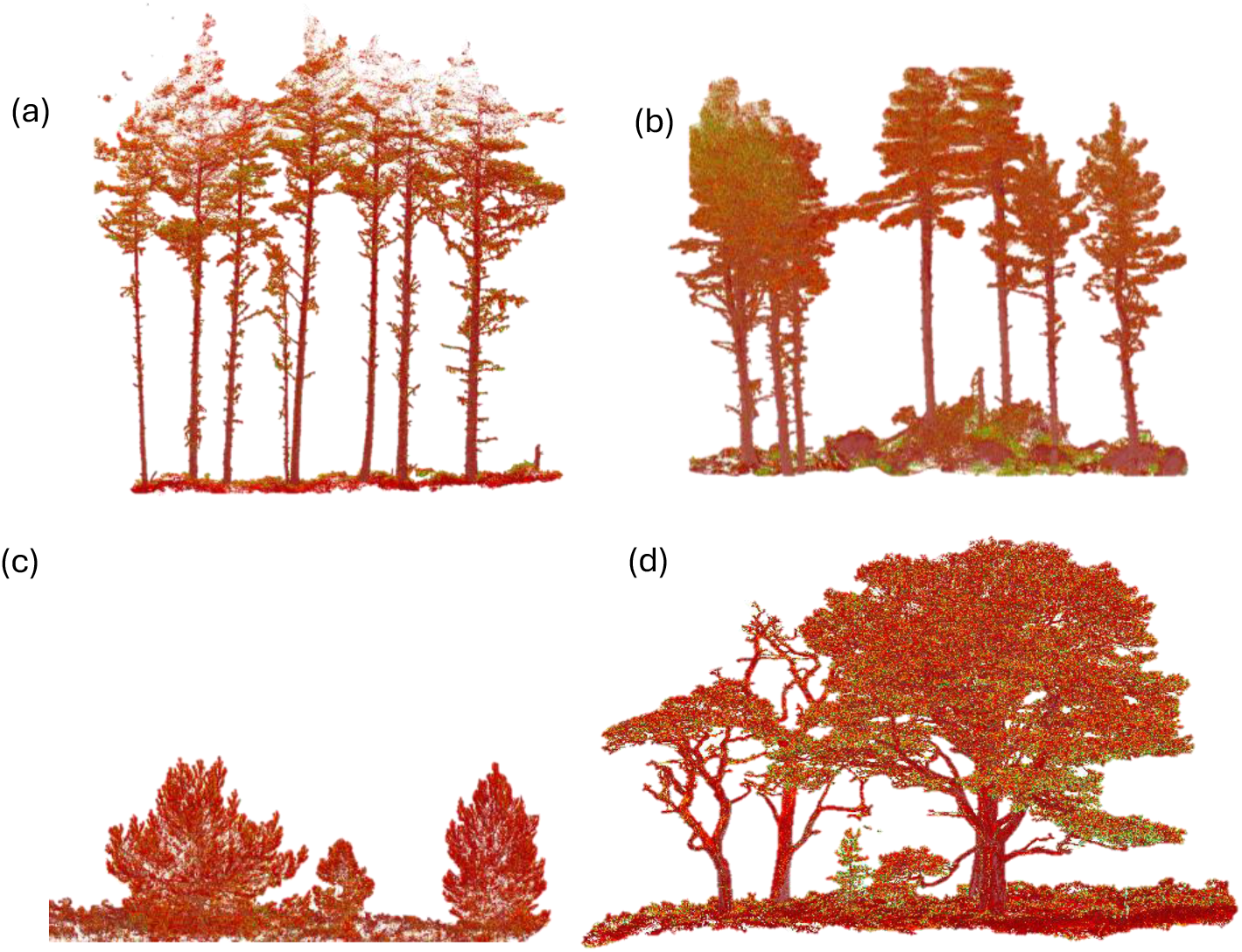
TLS point cloud examples from our four forest plot categories: (a) old plantation (OP), (b) old plantation managed (OPM), (c) natural regeneration (NR), (d) ancient woodland (AW).

AW was classified as areas that have been continuously wooded in Scotland since 1750 (NatureScot, 2023). AW plots typically contain trees of mixed ages, exhibit wide morphological variation, and are often located in relatively inaccessible areas, reducing the likelihood of extensive planting or management (Steven & Carlisle, 1959). Despite their ancient woodland status, AW plots suffered negative impacts through human influence over the course of their history, including high levels of deer grazing/browsing and historic felling pressures.

OP plots were occupied by trees of predominantly uniform age and height. The original high planting densities produce tall, slender trees with comparatively smaller diameter at breast height (DBH) and crown area relative to trees in AW. OP stands were also characterised by thin, dead branches extending up the trunks from self-pruning, reflecting intense competition for light beneath the canopy.

OPM plots underwent conservation management intervention by the Cairngorms Connect Partners within the past two-nine years. The management within these plots included deadwood creation via, ringbarking, winching and selective felling. This ‘plantation restructuring’ management aims to create canopy gaps with increased light availability, stimulating the development of a more complex, diverse understorey, promoting tree regeneration and enabling lateral expansion of tree crowns.

Finally, natural regeneration (NR) plots form part of the ‘regeneration zone’ of AW Inventory sites (NatureScot, 2023), falling in dispersal range of mature AW. They are characterised by extensive open shrub, and do not yet have a typical canopy layer. They are interspersed with young clusters of *P*. *sylvestris* and the occasional emergent tree.

All plots are dominated by *P. sylvestris,* but the understory does contain a scattering of regenerating saplings of other species: *Betula spp.*, *Populus tremula*, *Sorbus aucuparia*, *Juniperus communis*, and *Ilex aquifolium*. The shrub layer consists of *Calluna vulgaris, Juniperus communis, Vaccinium myrtillus* and *Vaccinium vitis-idaea* which grow amongst herbaceous plants, mosses and lichens.

### TLS data acquisition and pre-processing

We selected 32 plots, 30 x 30m, across the Cairngorms Connect area representing our different forest categories: eight AW, seven OP, ten OPM and seven NR (Fig. 1). We scanned plots in July 2024 and August 2025 using a Riegl VZ400i TLS (Riegl Laser Measurement Systems, Horn, Austria). We used a scan rate of 600 MHz and an angular resolution of 0.04 mrad under low wind conditions (gusts <12.00 mph), with no visible tree movement. We also avoided scanning in rain or fog to reduce noise caused by light refraction through precipitation. To minimise disturbance to Capercaillie (*Tetrao urogallus*; a Schedule 1 breeding bird) we avoided scanning outside the hours of 09:00 and 17:00 and during the core breeding season (April-June).

We followed a standard grid-based acquisition strategy, of approximately 10m spacing (Wilkes *et al*., 2017), resulting in a minimum of 16 scan positions per plot. At each scan position, we collected two scans: an upright scan and a tilt scan, in which we oriented the scanner 90° horizontally to enhance canopy coverage. This is considered standard practice for the Riegl VZ400i, which does not have a full hemispherical field of view, Wilkes *et al*., 2017. When dense vegetation limited line of sight to the tree stems, we added additional scans to maximise coverage and minimise occlusion. We placed cylindrical high-reflective targets to support post-processing and maintained real-time location corrections to the TLS using an Emlid Reach RS3 GNSS base station.

We co-registered the individual scans in RiSCAN PRO (Reigl, Horn, Austria) and generated full stand-level point clouds using the rxp-pipeline (Wilkes, 2023). We cropped the point clouds to include a 2.5m convex hull buffer around the scan positions, down sampled to point density of 1cm, and filtered by reflectance (<15dB) and point deviation (>15). We manually removed outlier points along the z-axis that were visually above the canopy using CloudCompare v2.13.1 (*CloudCompare*, 2024).

We pre-processed the point clouds in R (R Core Team, 2025) using the *lidR* package (Roussel et al., 2020). We created digital terrain models (DTM) for each plot using the cloth simulation filter (CSF; Zhang et al., 2016) with default parameters. We normalised the plot heights by subtracting ground elevation from each point. Given the typically low stature of Cairngorm pinewoods understory, we only removed points within 2 cm of the normalised ground (0 m), rather than a fully classified ground layer, as the understory holds high ecological value and contributes significantly to plot structure. We generated CHMs using the pit-free algorithm implemented in *lidR* (Khosravipour et al., 2014; Roussel et al., 2020), with a sub circle diameter equal to the resolution value. For voxel creation we generated a 3D grid at each specified resolution using a custom voxel indexing to the extent of each point clouds bounding box.

### Metric choice and resolution sampling

We selected 21 FSC metrics that have been used in the literature to characterise FSC in relation to forest management or biodiversity (Table 1 and Table S1). We categorised these metrics, by adapting Feigl *et al.,* (2025) metric grouping, into five groups: density, fractal, height, openness and rugosity. Of these metrics, 20 require one or more user-defined resolution parameter: (1) voxel size, (2) CHM pixel resolution, and (3) vertical height bin size. The final metric, box dimension, did not require a voxel or CHM resolution choice (Seidel, 2018).

For each of the 20 resolution-dependent metrics, we calculated 31 values across all 32 plots using either CHM, voxel or height bin resolution ranging from 0.02m to 1.5 m in 0.05m increments. We chose this range to capture values used in existing TLS or MLS based FSC studies but also extended to larger values to align with resolutions more frequently used for airborne and drone lidar studies (Supporting Information Table S3).

We selected eight metrics that require a voxel-size resolution choice (metrics 1-8, Table 1): 3D gap fraction (gap_frac_3D), vertical space filling (SFz), Gini coefficient of vertical space filling (GINIz), Gini coefficient of horizontal space filling (GINIxy), coefficient of variation of vertical space filling (CVz), coefficient of variation of horizontal space filling (CVxy), coefficient of variation of leaf area density (CV_LAD) and Effective Number of Layers (ENL). A voxel was classified as filled if it contained one or more points. Foliage height diversity (FHD, 9) required both voxel-size and height-bin size resolution choices simultaneously, and Vertical Complexity Index (VCI, 10) required just height bin. Metrics 12 – 18 required a CHM pixel resolution choice: mean, max, standard deviation, coefficient of variation and Gini coefficient of height (*metric_name*_*height), rumple index (rumple) and rugosity (rugosity). Metrics 19 – 21: gap fraction (gap_frac_33), canopy cover (cover_33) and coefficient of variation of gap area (CV_gap_area_33), also required CHM pixel-resolution but were calculated using a threshold of 33% of canopy height, applying relative rather than absolute height thresholds to facilitate comparison among plots of differing canopy height (Silva et al., 2019; Willim et al., 2020).

### Forest structural category rank stability to metric resolution value changes

We established forest category complexity ranking at each resolution gradient for each metric using the mean result of each metric at each resolution size. This produced a ranking sequence for each metric. From each ranking sequence we could identify any changes in ranking order of forest categories and at which resolution value this change happened. To simplify our interpretation, we binned the resolution gradient into fine (<0.1m), small (0.1m – 0.5m), medium (0.5m-1m) and coarse (>1m) windows. We identified metrics as stable if they had ≤1 change in ranking order across the entire gradient. We allowed moderation of one change in ranking order as this would still allow for a window of stability within the gradient. Any metrics with >1 change were deemed unstable and discounted from further analysis.

### Selecting resolution size for maximising forest category discernability

We defined the optimal resolution for each metric as the resolution at which the greatest separation between forest categories was observed. We summed the absolute differences between the metric value means of each category at each resolution value and selected the value with the largest summed difference. Chosen resolution values for each stable metric are shown in Table 2

**Table 2.** Resolution values chosen for further analysis after completing ranking analysis. Values were chosen as the resolution value at which the separation between mean metric values across forest categories was the greatest, using absolute pairwise mean category differences. CHM=canopy height model.

| FSC Metric | Group | Resolution choice |  |
| --- | --- | --- | --- |
|  |  | type | Value |
| Gap fraction 2D | Openness | CHM resolution | 0.02 |
| Gini height | Rugosity | CHM resolution | 0.02 |
| CV height | Rugosity | CHM resolution | 0.05 |
| Canopy cover | Openness | CHM resolution | 0.05 |
| SD height | Rugosity | CHM resolution | 0.10 |
| SFz | Density | Voxel size | 0.20 |
| Max height | Height | CHM resolution | 0.20 |
| CV LAD | Density | Voxel size | 0.70 |
| VCI | Density | Voxel size | 1.45 |
| Gap fraction 3D | Density | Voxel size | 1.45 |
| Mean height | Height | CHM resolution | 1.45 |

## Results

### FSC metric resolution choice influences forest category structural comlexity rankings

Six metrics ranked forest categories consistently, regardless of resolution value (*Gini height, CV height, VCI, SFz, gap frac 3D* and *CV LAD*). There were a total of 52 changes in rank order for the remaining 14 metrics across the resolution value range. Of these, 9.6% rank changes occurred at fine-scale, 30.8% at small-scale, 32.7% were at medium-scale and 26.9% were at coarse-scale. CHM-based metrics (n= 10) displayed a total of 28 rank changes and voxel/height bin metrics (n=10) exhibited the remaining 24. Voxel-based metrics typically demonstrated higher ranking instability at fine and small-scale resolution values (26.0% of total changes) compared with CHM-based metrics (13.5%; see Fig. 3b). In contrast, CHM-based metrics showed relative instability at medium-scale and coarse-scale (39.6%).

**Fig. 3.**
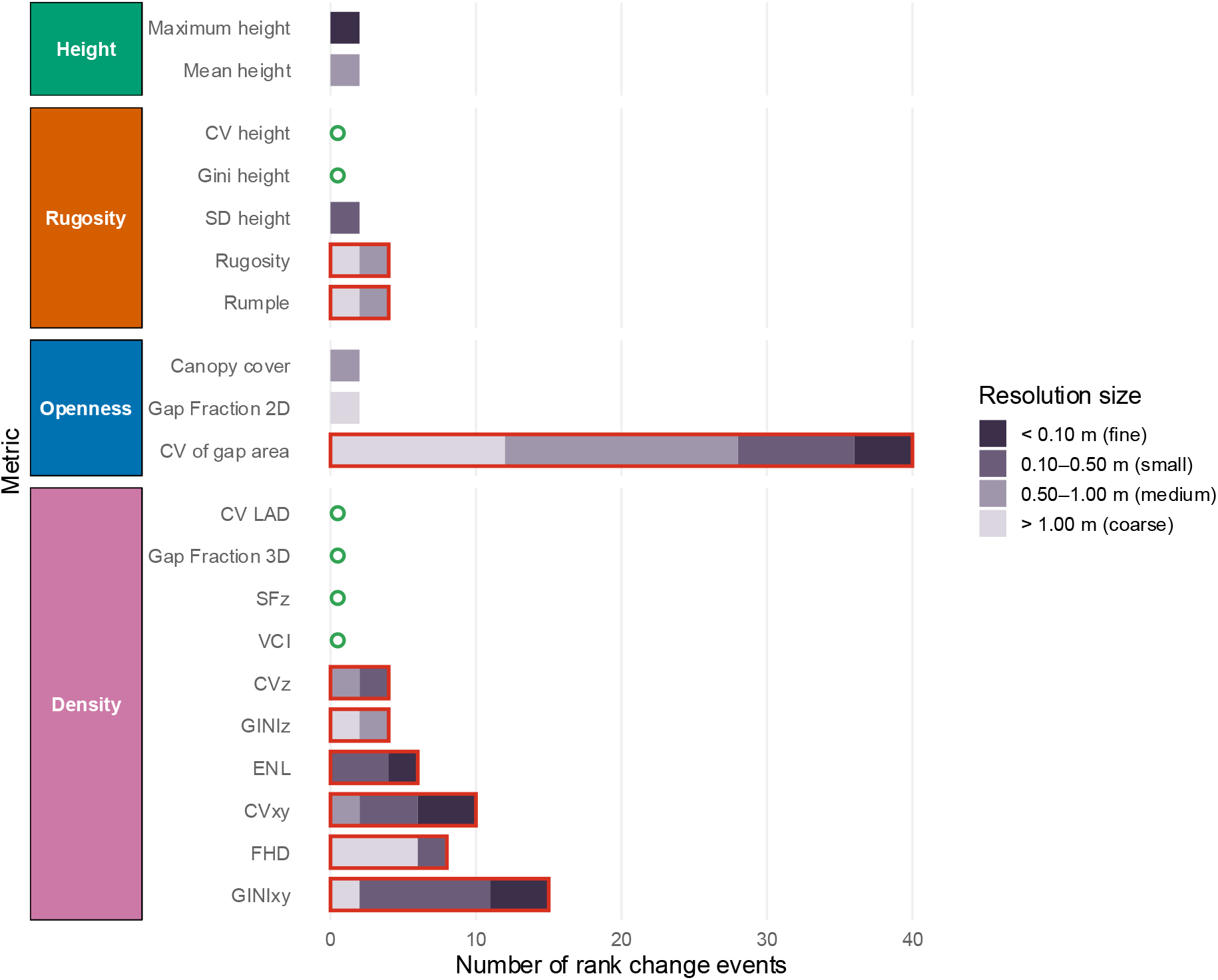
Bar charts showing the number of category rank change events for each metric over the range of resolutions tested, with metrics grouped by calculation method (see Feigl *et al.,* 2025). Metric bars are stacked by colour to indicate the number of rank change events at different resolution sizes: rank change events that occurred < 0.10m i.e. fine scale resolution/voxel size (dark grey); rank change event occurred between 0.10-0.50m (medium grey); rank change events between 0.50-1.00m (grey) and rank change events above 1m resolution (light grey). Metric bars outlined in red indicate metrics that were considered unstable with >1 rank change.

All methodological groupings contained at least one metric with no or only one rank change, but metric stability across resolution change varied by methodological grouping and resolution type used (Fig. 3a). Five out of the 14 metrics had only one rank order change and were carried forward for agreement analysis along with the six stable metrics: *mean height* and *max height* (height metrics); *CV LAD, SFz, VCI, gap frac 3D* (density metrics); *gap frac 2D*, *canopy cover* (openness metrics) and; CV_height, *sd height* and *Gini height* (rugosity metrics). Nine metrics were discounted as unstable: rumple, rugosity (rugosity metrics); *CV gap area* (openness metrics); *GINIz, GINIxy, FHD, ENL, CVz* and *CVxy* (density metrics).

### CHM pixel resolution influenced forest category ranking in four out of ten metrics

Openness metrics, *gap fraction 2D* and *canopy cover,* had one rank change each and were both considered stable at small or fine-scale resolution values (0.02-0.1m and 0.1-0.5m), with each rank change occurring at medium or coarse-scales. However, *CV gap area* (20 rank changes) was highly unstable across the range of resolutions.

Both height group metrics, *max* and *mean height*, had one rank change each and were considered comparatively stable. *Max height* had single rank change at 0.05m and no following changes; *mean height* had one change at 0.9m but ranking remained stable at fine, small and coarse-scale resolution values.

In the rugosity group, *CV height* and *Gini height* were clearly stable, with no rank changes across the full resolution range, but others showed varied stability. *Sd height* showed some stability, with only one rank change at 0.35m and otherwise stable rankings at fine, medium and coarse resolution values. *Rugosity* and *rumple index* both had two rank changes after 0.50m and were discounted.

*Voxel and height bin* value *influenced forest category ranking in six out of ten metrics*

Four metrics in the density group exhibited no rank changes, *VCI, SFz, gap frac 3D and CV LAD,* and were classified as stable. Rank changes were most frequent at small-scale (11 rank changes from 0.1-0.5m). *FHD* had four rank change at 0.05m, 1.3m, 1.4m and 1.45m, and *CVz* had two at 0.3m and 0.9m. Both metrics were discounted. *GINIz* had a total of four rank change events above 0.50m and was also discounted.

The least stable density metrics were *CVxy* (four rank changes) and *ENL* (five rank changes), with rank changes at both fine and small-scale resolution values. *ENL* exhibited a cluster of five rank-changes between voxel values 0.05 – 0.35m; this range includes the standard voxel size used in the literature of 0.20m, highlighting the challenge of reliable usage.

### Metrics can distinguish NR from other categories but OP and OPM are statistically indistinguishable

Based on patterns in Fig. 4 we placed metrics into three concordant groups defined by agreed ranking order of forest categories with other metrics, and one non-concordant group. Here, we also include box dimension as a holistic, resolution-choice free metric used in the literature. Our four groups (Fig. 4) are as follows. Group 1 ranks forest categories as OP > OPM > AW >NR, and these metrics primarily capture canopy cover and vertical heterogeneity: *VCI, SFz, BoxD, canopy cover.* Group 2 metrics had an opposite ranking order to group 1 (NR > AW > OPM > OP) and included gap and height inequality-based metrics, consisting of: *Gini height, CV height, gap fraction 2D* and *gap fraction 3D*. Group 3 metrics ranked plots as OPM > OP > AW > NR, included height metrics: *max height* and *mean height. CV LAD* and *SD height* ranked differently to these groups (NR > OP > OPM > AW and OPM > AW > OP > NR, respectively) and so are presented in a group of non-concordance. NR has the highest complexity ranking in group 2 (Fig. 4) which contains height inequality and openness metrics. In contrast, group 1 and group 3, which incorporate density, openness, fractal and height metrics rank NR with the lowest complexity.

**Fig. 4.**
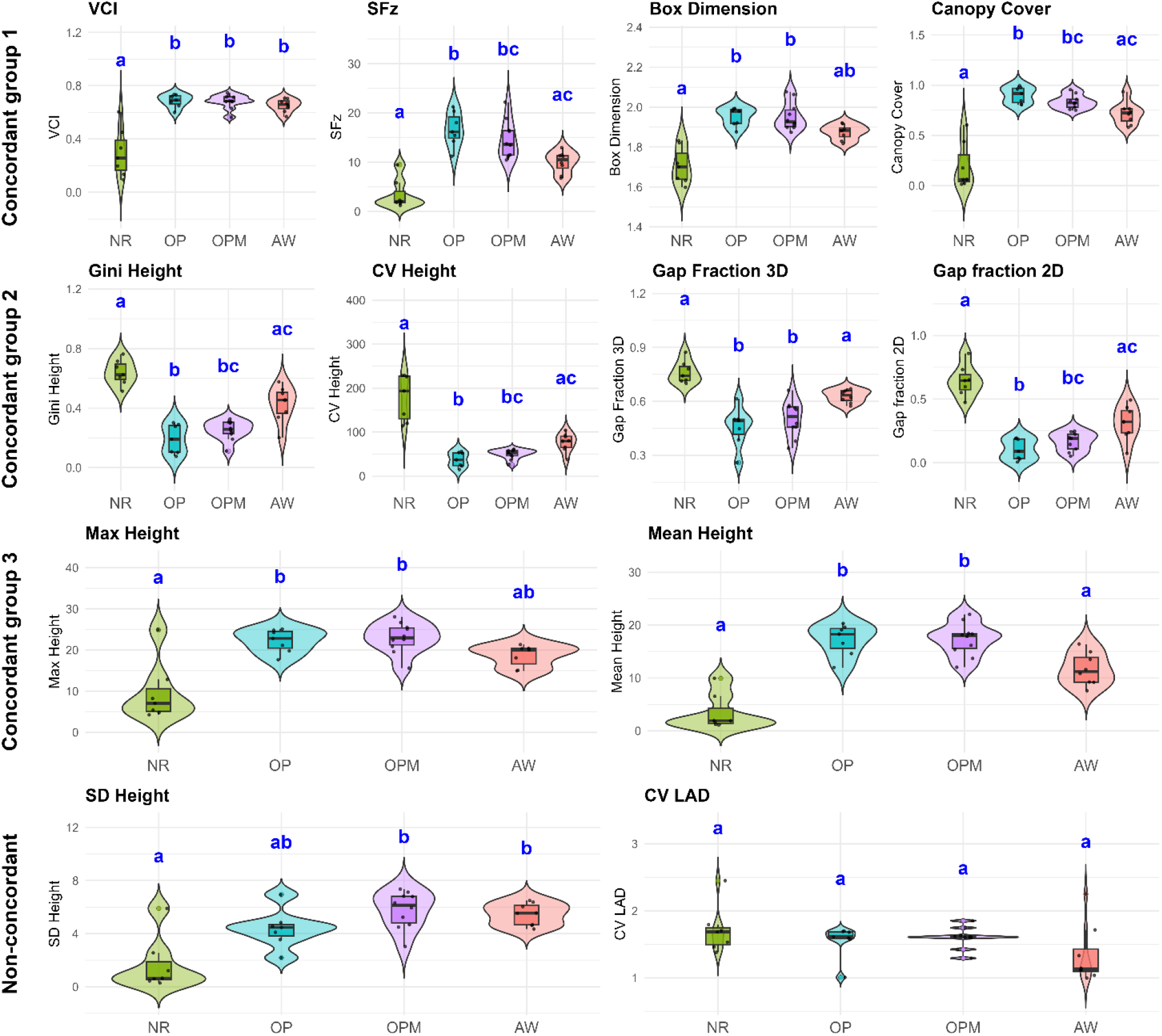
Discernability analysis. Violin plots of metric results, showing ranking order of forest categories. Letters represent statistical differences as per a Kruskal-Wallis non-parametric test with a Dunn BH-adjusted post hoc test. Metrics are grouped by concordance, defined by the agreement of ranking order of forest categories.

A non-parametric Kruskal-Wallis (Kruskal & Wallis, 1952) test showed that all metrics found statistical differences between at least two forest categories, except for *CV LAD,* which was the only metric unable to significantly distinguish any forest categories (H₃ = 4.63, p = 0.2). Post-hoc pairwise comparisons, showed that this was primarily driven by strong separation between NR and the other taller forest categories, but that AW, OP and OPM often shared similar FSC values (Dunn test, BH-adjusted P < 0.05 (Benjamini & Hochberg, 1995; Dunn, 1964)). There was no significant distinction between OP and OPM by any metric, indicating a structural overlap between these forest categories. This overlap is evident in Figs. 4 and 5.

**Fig. 5.**
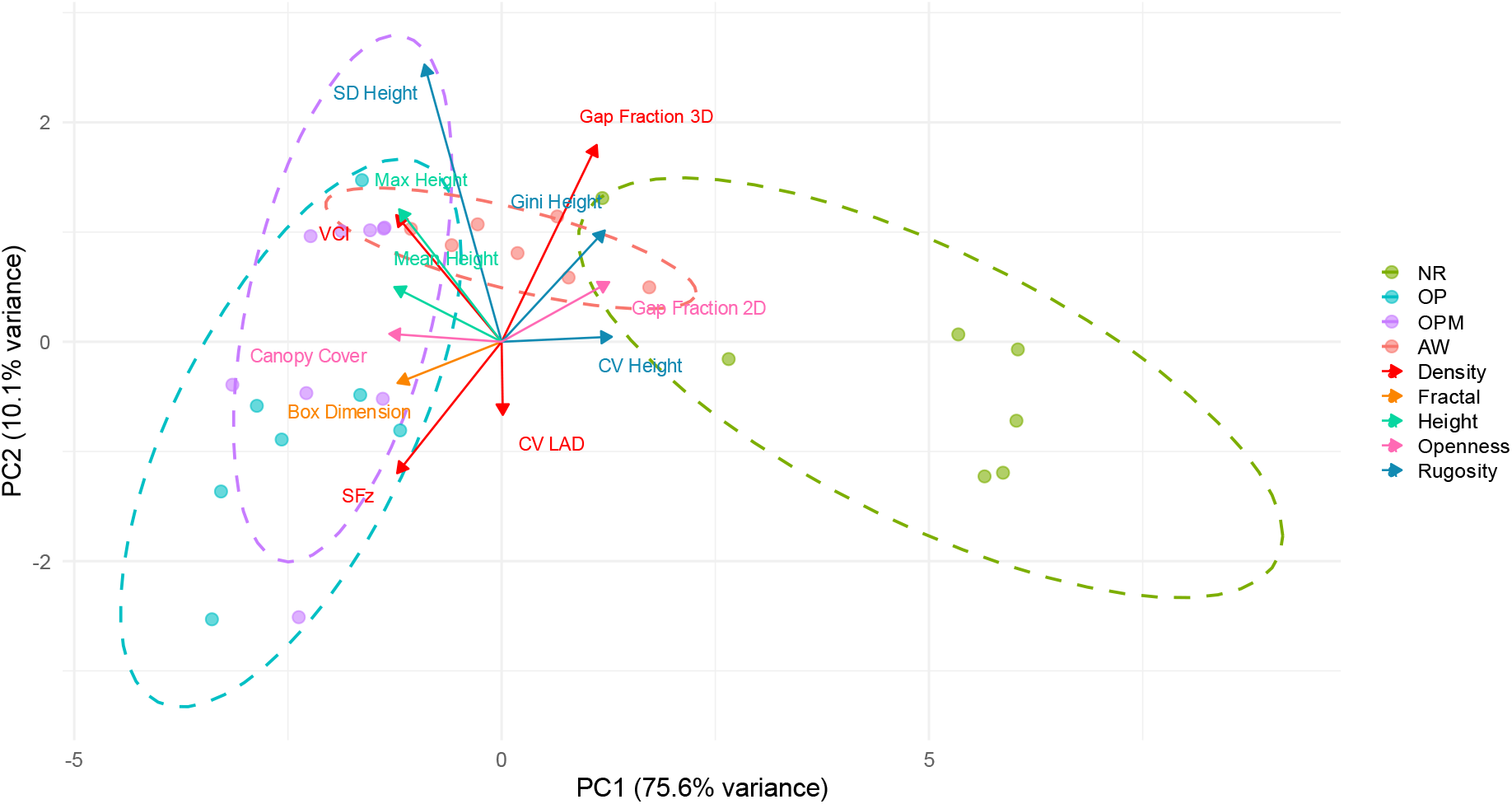
A biplot principal component analysis of structural complexity metrics across four categories of forest structure: (NR) natural regeneration, (OP) old plantation, (OPM), old plantation managed for conservation and (AW), ancient woodland. Coloured points represent individual plots of each category and dashed ellipsoids represent 95% confidence regions around each group. The arrows show variable loadings on PC1 and PC2, coloured by metric group (density, fractal, height, openness and rugosity). PC1 (75.6% variance) separates NR from the remaining groups, driven by gap fraction and rugosity metrics. PC2 (10.1%) captures secondary variation among OP, OPM and AW driven by mean height and gap fraction of the herb layer.

AW was not always distinguished from OP, or even sometimes OPM, by FSC metrics. *Gap fraction 2D* (H₃ = 21.92, P <0.05)*, CV height* (H₃ = 23.64, P <0.05)*, Gini height* (H₃ = 22.87, P <0.05)*, Canopy cover* (H₃ = 21.47, P <0.05), and *SFz* (H₃ = 21.92, P <0.05), significantly discriminated AW from OP but not OPM (Dunn test, BH-adjusted P < 0.05). However, despite significant overall test effects; *VCI* (H₃ = 16.7, P <0.05)*, Box dimension* (H₃ = 20.34, P <0.05)*, sd height* (H₃ = 13.95, P <0.05), and *max height* (H₃ = 13.7, P <0.05), in pairwise analysis these metrics did not significantly distinguish AW from OP and OPM. *Gap fraction 3D* (H₃ = 23.11, P <0.05) and *mean height* (H₃ = 21.88, P <0.05) were the only metrics to significantly differentiate AW from OP and OPM categories in pairwise comparisons but in each case, FSC of AW was not significantly different from NR.

### Canopy cover and high CV height drive NR separation

To further understand similarities and differences in rankings across FSC metrics and forest categories, we performed a PCA on all metrics assessed as stable and evaluated scores weights and groupings (Fig. 5). PC1 captures 75.6% of the variance across our plots and mostly separates NR plots from OP, OPM and AW categories. Metrics related to openness and height inequality (*CV height* and *canopy cover*) drive the group separation along PC1. NR plots are widely dispersed with high values of PC1, indicating high structural variability within this category, whereas OP, OPM and AW cluster with greater within-group consistency, with OP and OPM having lower PC1 values and AW having medium values. *SFz and SD* height separate OP and OPM, and AW occupies an intermediate space characterised by high gap fraction and height.

PC2 (10.1% variance) captures the secondary variation among the taller forest categories. OPM plots had the highest PC2 values, influenced strongly by *sd height*, suggesting greater canopy height heterogeneity relative to OP forests, despite the metrics’ inability to statistically differentiate these. OP generally had negative PC2 values, driven by higher *SFz,* OP and OPM overlap for both PC1 and PC2, suggesting they are less distinctive with greater within-category variance than AW, despite visual assessment of differences being clear in the field (Figs. 2 and S1). AW occupies a central location along PC1 and a positive position on PC2.

## Discussion

Forest structural complexity is increasingly used as a proxy for biodiversity in conservation, restoration and forest management (Supporting Information Table S1), making the reliability of lidar-derived structural metrics a critical issue. Here we evaluated the stability, discernability and ecological interpretability of 21 TLS-derived forest structural complexity metrics across structurally contrasting Scots pine forests, identifying several challenges that limit their ecological application.

Our ranking analysis (Fig. 3) has direct implications for metrics already widespread in ecology. Resolution selection influenced not only metric values, but also the ecological conclusions drawn from them (Figs. 3 and 4). Altering resolution size of metrics changed the structural ranking order of forest categories but this varied by metric, metric grouping and resolution parameter type. Among these metrics, Effective Number of Layers (ENL) has become one of the most widely adopted predictors of biodiversity (Cordeiro Pereira et al., 2026; Rappa et al., 2023; Wildermuth et al., 2023). Most studies adopt the original 0.2 m voxel size proposed by Ehbrecht et al. (2016), often without explicitly evaluating whether this parameterisation is appropriate, despite the original authors noting that ENL is sensitive to voxel resolution. We observed forest category rankings change five times for ENL between 0.02–0.5m, suggesting that no voxel size within this range can be assumed to provide reliable ecological inference without prior testing. This does not, however, invalidate the results from the latter studies as the resulting correlations remain internally consistent within each study system. Rather, it limits the transferability and reproducibility of ENL as an ecological indicator across study systems, complicating comparisons among studies and future meta-analyses. We recommend that future applications of forest structural complexity metrics should therefore demonstrate that the selected voxel size is reliable within the specified study system. Sensitivity analyses should become routine before structural metrics are adopted as predictors of biodiversity or restoration outcomes, with explicit justification of the chosen parameterisation.

Our results reinforce a common view that FSC is inherently multidimensional, such that reliance on a single lidar-derived metric may overlook ecologically meaningful dimensions of forest structure (Atkins et al., 2023b; LaRue et al., 2023; Rosen et al., 2024). Box dimension, a holistic metric requiring fewer parameter selections than voxel-based alternatives, did not capture height inequality and gap fraction which are characteristic of both NR and AW forest plots, nor did it provide discriminatory power between OP and OPM, contributing to their overlap in Fig. 5. It did, however, produce category rankings similar to VCI, SFz and canopy cover metrics, suggesting that box dimension primarily captures vegetation occupancy rather than integrating multiple dimensions of forest structure. Plot geometry may partly explain this pattern. Reuter et al. (2026) note that box dimension decreases as plot side length increases relative to canopy height. Our plots maintained a consistent width whilst canopy height differed, this relationship may have influenced metric performance. Other holistic measures of structural complexity from the literature also appear unlikely to fully overcome these limitations. For example, Canopy Entropy (CE) (Liu et al., 2022) is strongly correlated with Box Dimension (Reuter et al., 2026), but its calculation requires selection of a KDE bandwidth, with Liu et al. (2022) selecting a value of 0.2 m through trial and error, noting that bandwidth influences the smoothness of the estimated probability density and therefore the resulting CE estimates. Likewise, the Stand Structural Complexity Index (Ehbrecht et al., 2017) was developed for single-scan TLS and is therefore not directly applicable to our multi-scan dataset (Liu et al., 2022). Collectively, these findings suggest that no currently available holistic metric fully captures the multiple structural dimensions that distinguish contrasting forest conditions, reinforcing the need for complementary metrics rather than reliance on a single measure of forest structural complexity.

The need for reliable structural metrics extends beyond individual ecological studies. National terrestrial, mobile, airborne lidar campaigns, emerging spaceborne products and associated initiatives such as Defra’s ReForeSt project, are producing operational maps of forest structural diversity to support biodiversity assessment and woodland management (Broome et al., n.d.; de Conto et al., 2024; Holvoet et al., 2025; Scottish Government, 2026). In these products, the processing decisions such as those explored here are made upstream and are largely invisible to the end user. Although our results do not transfer quantitatively to other lidar platforms, the underlying principle remains the same: ecological inference from complexity metrics depends on parameterisation. As these metrics become embedded within landscape-scale monitoring and management, the robustness and ecological interpretability of their underlying metrics should be explicitly evaluated, particularly across airborne and spaceborne lidar systems.

We evaluated whether lidar-derived structural complexity metrics could detect the early effects of restructuring management implemented by the Cairngorms Connect project. Although no metric detected a statistically significant difference between old plantation (OP) and old plantation managed with re-structuring (OPM), several of our analyses suggested that restructuring effects are beginning to emerge in metric estimates (Fig. 4). OPM consistently ranked higher than OP for openness and height inequality metrics (concordant group 2) and lower for canopy-filling metrics (group 1). Likewise, the limited separation between OP and OPM observed in the PCA was driven primarily by standard deviation of height and vertical space filling (SFz), consistent with early crown expansion into created gaps and the increasing canopy height heterogeneity that is expected following restructuring (Cheng et al., 2025; Saarinen et al., 2021). Standard deviation of height and SFz therefore emerge as promising candidate indicators for longitudinal monitoring of restructuring management and warrant further evaluation. Several additional metrics (canopy cover, Gini height, CV height and gap fraction 2D) distinguished ancient woodland (AW) from OP while remaining indistinguishable from OPM. Restructuring management aims to promote structural characteristics associated with ancient woodland, it could follow that these metrics may be particularly sensitive to the early stages of this transition. Given that management was completed two-nine years prior to surveying, whereas AW structure has developed over centuries, a substantial convergence from OPM to AW would not be expected (Constantz et al., 2025; Kuuluvainen, 2009). Instead, our results suggest that lidar-derived structural metrics can detect subtle early changes associated with restructuring management, but that the ecological interpretation depends strongly on the metric selected. Importantly, the apparent similarity between OPM and AW reflects a non-significant statistical difference and should therefore be interpreted as suggestive rather than conclusive. Together, these results suggest that metrics describing canopy openness and vertical heterogeneity may detect the earliest structural responses to restructuring management, whereas metrics describing overall vegetation occupancy respond over longer timescales.

Contrary to our hypothesis, AW was not consistently identified as the structurally most complex forest type and instead occupied an intermediate ranking between NR and OP/OPM (Figs. 4 and 5). This intermediate position reflects the coexistence of structural characteristics shared with each forest category, rather than reduced structural complexity. Metrics describing vegetation occupancy grouped AW with OP and OPM, reflecting the persistence of mature, well-developed canopy and taller trees. In contrast, metrics describing canopy openness and height inequality grouped AW closer to NR. AW serves less as an endpoint reference but more readily provides supporting evidence that structural complexity is multidimensional, as what distinguishes AW is not high value on any single metric but is simultaneously expressed by moderate canopy occupancy, openness and height heterogeneity; a combination of dimensions that no single one of our individual metrics could capture alone.

We deliberately standardised data acquisition to isolate differences among structural metrics. We restricted analysis to stands of single species to reduce the confounding influence of tree species composition on structural complexity (Coverdale & Davies, 2023); we used a standardised rigorous TLS sampling strategy which limited occlusion bias and subsampled each plot to 1cm for across plot consistency. These choices allowed us to evaluate metrics under controlled conditions, but these controls also define the boundaries of our conclusion. The patterns observed here may differ if derived from alternative lidar platforms (e.g. ALS, ULS or MLS), survey strategies or point cloud densities which are known to influence metric results (Cimdins et al., 2025; Donager et al., 2018; LaRue et al., 2022). Further work should support this emerging theme to examine whether complexity metrics remain stable when *both* resolution choice and data acquisition techniques vary.

In conclusion, our research shows that resolution choice does not merely alter values derived from forest structural complexity metrics but can lead to contrasting ecological interpretations of our data. ENL is a popular structural complexity metric, yet we found it to be unstable, when tested in the context of our study system. We identified metrics that produced stable ecological inference, while recognising that their performance may vary among sensors and ecosystems. We recommend that resolution-metric sensitivity analysis become a routine component of lidar based biodiversity studies and that parameter selection is explicitly justified. Stable structural complexity metrics clearly distinguished contrasting forest types but showed minimal difference between recent management practice changes. Consequently, we suggest that managers wishing to evaluate their re-structuring management using similar metrics to do so with a repeated-plot, longitudinal study design and realistic expectations of timescale, rather than comparing to a baseline category. More broadly, as lidar-derived structural metrics transition from research tools to regional, national and global indicators, confidence in their stability and ecological interpretability becomes increasingly important. Structural change is a dynamic, multi-generational process that is unique to each ecological system and should be evaluated as such. Ensuring that structural complexity metrics provide reproducible ecological inference will be essential if they are to support evidence-based conservation policy and management.

## Data availability statement

TLS point clouds will be made openly available on Zenodo upon acceptance of the manuscript.

For the purposes of review, data is available here:

https://zenodo.org/records/22801247?token=eyJhbGciOiJIUzUxMiJ9.eyJpZCI6ImU1OGIwYzkwLTI3NjgtNDVjMy1iYzM0LWRiZmQ0MmY5NDJjMyIsImRhdGEiOnt9LCJyYW5kb20iOiJhNDlkY2ZlNzY1YzJiYmJmMDBlMDg2NjZlMTQ4YTIwYSJ9.iIWRXUZ-bCNORkf8Sz-D0ec87BWGO5mk4jQk4hvnFdjmtLoKl3okZshEpvVphAXwg_3IuTTTcGSDnuOElqLbdg

## References

Almeida, D. R. A., Stark, S., Silva, C., Hamamura, C., & Valbuena, R. (2021). _leafR: Calculates the Leaf Area Index (LAD) and Other Related Functions_. (Version 0.3.5) [Computer software]. 10.32614/CRAN.package.leafR

Amphlett, A. (2022). A review of the vascular plant flora of the Cairngorms Connect project area, Scotland, and some possible implications of forest expansion to the natural tree line. British & Irish Botany, 4. 10.33928/bib.2022.04.166

Anderson, C. T., Dietz, S. L., Pokswinski, S. M., Jenkins, A. M., Kaeser, M. J., Hiers, J. K., & Pelc, B. D. (2021). Traditional field metrics and terrestrial LiDAR predict plant richness in southern pine forests. Forest Ecology and Management, 491, 119118. 10.1016/j.foreco.2021.119118

Asbeck, T., Benneter, A., Huber, A., Margaritis, D., Buse, J., Popa, F., Pyttel, P., Förschler, M., Gärtner, S., & Bauhus, J. (2023). Enhancing structural complexity: An experiment conducted in the Black Forest National Park, Germany. Ecology and Evolution, 13(1), e9732. 10.1002/ece3.9732

Atkins, J. W., Bhatt, P., Carrasco, L., Francis, E., Garabedian, J. E., Hakkenberg, C. R., Hardiman, B. S., Jung, J., Koirala, A., LaRue, E. A., Oh, S., Shao, G., Shao, G., Shugart, H. H., Spiers, A., Stovall, A. E. L., Surasinghe, T. D., Tai, X., Zhai, L., … Krause, K. (2023). Integrating forest structural diversity measurement into ecological research. Ecosphere, 14(9), e4633. 10.1002/ecs2.4633

Atkins, J. W., Costanza, J., Dahlin, K. M., Dannenberg, M. P., Elmore, A. J., Fitzpatrick, M. C., Hakkenberg, C. R., Hardiman, B. S., Kamoske, A., LaRue, E. A., Silva, C. A., Stovall, A. E. L., & Tielens, E. K. (2023). Scale dependency of lidar-derived forest structural diversity. Methods in Ecology and Evolution, 14(2), 708–723. 10.1111/2041-210X.14040

Béland, M., & Baldocchi, D. (2020). Is foliage clumping an outcome of resource limitations within forests? Agricultural and Forest Meteorology, 295, 108185. 10.1016/j.agrformet.2020.108185

Benjamini, Y., & Hochberg, Y. (1995). Controlling the False Discovery Rate: A Practical and Powerful Approach to Multiple Testing. Journal of the Royal Statistical Society: Series B (Methodological), 57(1), 289–300. 10.1111/j.2517-6161.1995.tb02031.x

Blakey, R. V., Law, B. S., Kingsford, R. T., & Stoklosa, J. (2017). Terrestrial laser scanning reveals below-canopy bat trait relationships with forest structure. Remote Sensing of Environment, 198, 40–51. 10.1016/j.rse.2017.05.038

Bouvier, M., Durrieu, S., Fournier, R. A., & Renaud, J.-P. (2015). Generalizing predictive models of forest inventory attributes using an area-based approach with airborne LiDAR data. Remote Sensing of Environment, 156, 322–334. 10.1016/j.rse.2014.10.004

Broome, A., Scopes, E., Whittacker, C., Watts, K., & Hughes, S. (n.d.). ReForeSt: Remote sensing for Forest management and Structural Diversity. Forest Research. Retrieved 6 August 2026, from https://www.forestresearch.gov.uk/research/reforest-remote-sensing-for-forest-management-and-structural-diversity/

Cairngorms Connect. (n.d.). Cairngorms Connect. Retrieved 15 July 2026, from https://cairngormsconnect.org.uk/

Calders, K., Adams, J., Armston, J., Bartholomeus, H., Bauwens, S., Bentley, L. P., Chave, J., Danson, F. M., Demol, M., Disney, M., Gaulton, R., Krishna Moorthy, S. M., Levick, S. R., Saarinen, N., Schaaf, C., Stovall, A., Terryn, L., Wilkes, P., & Verbeeck, H. (2020). Terrestrial laser scanning in forest ecology: Expanding the horizon. Remote Sensing of Environment, 251, 112102. 10.1016/j.rse.2020.112102

Carrasco, L., Giam, X., Papes, M., & Sheldon, K. (2019). Metrics of Lidar-Derived 3D Vegetation Structure Reveal Contrasting Effects of Horizontal and Vertical Forest Heterogeneity on Bird Species Richness. Remote Sensing, 11(7), 743. 10.3390/rs11070743

Chen, Y., Liu, J., Yao, X., Deng, Y., Hao, Z., Lin, L., Wu, N., & Yu, K. (2022). Effect of layer thickness and voxel size inversion on leaf area density based on the voxel-based canopy profiling method. Journal of Forestry Research, 33(5), 1545–1558. 10.1007/s11676-021-01440-7

Cheng, F., Zhao, X., Luo, Y., Tian, J., Sun, Y., Wang, H., Wang, Q., Zhu, J., Yang, C., He, H., & Liu, G. (2025). Effects of thinning on canopy structure, forest productivity, and productivity stability in mixed conifer-broadleaf forest: Insights from a LiDAR survey. Journal of Environmental Management, 385, 125707. 10.1016/j.jenvman.2025.125707

Cifuentes, R., Van der Zande, D., Farifteh, J., Salas, C., & Coppin, P. (2014). Effects of voxel size and sampling setup on the estimation of forest canopy gap fraction from terrestrial laser scanning data. Agricultural and Forest Meteorology, 194, 230–240. 10.1016/j.agrformet.2014.04.013

Cimdins, R., Yrttimaa, T., Hyyppä, J., Vastaranta, M., & Kankare, V. (2025). Capturing trends in forest structural complexity development using laser scanning techniques. *Trees*, Forests and People, 21, 100954. 10.1016/j.tfp.2025.100954

CloudCompare (Version 2.13.1). (2024). [Computer software]. https://www.cloudcompare.org/

Constantz, B. M., Stella, J. C., & Holl, K. D. (2025). Diverging restoration pathways for overstory and understory communities in a Mediterranean-climate riparian ecosystem. Ecological Applications, 35(3), e70043. 10.1002/eap.70043

Cordeiro Pereira, J. M., Klingenfuß, S., Basile, M., Frey, J., Mikusiński, G., & Storch, I. (2026). Disentangling the effects of multifunctional forestry practices on the abundances of birds and their invertebrate prey. Ecological Applications, 36(2), e70198. 10.1002/eap.70198

Coverdale, T. C., & Davies, A. B. (2023). Unravelling the relationship between plant diversity and vegetation structural complexity: A review and theoretical framework. Journal of Ecology, 111(7), 1378–1395. 10.1111/1365-2745.14068

de Conto, T., Armston, J., & Dubayah, R. (2024). Characterizing the structural complexity of the Earth’s forests with spaceborne lidar. Nature Communications, 15(1), 8116. 10.1038/s41467-024-52468-2

Deere, N. J., Guillera-Arroita, G., Swinfield, T., Milodowski, D. T., Coomes, D. A., Bernard, H., Reynolds, G., Davies, Z. G., & Struebig, M. J. (2020). Maximizing the value of forest restoration for tropical mammals by detecting three-dimensional habitat associations. Proceedings of the National Academy of Sciences, 117(42), 26254–26262. 10.1073/pnas.2001823117

Disney, M. (2019). Terrestrial LiDAR: A three-dimensional revolution in how we look at trees. New Phytologist, 222(4), 1736–1741. 10.1111/nph.15517

Donager, J. J., Sankey, T. Ts., Sankey, J. B., Sanchez Meador, A. J., Springer, A. E., & Bailey, J. D. (2018). Examining Forest Structure With Terrestrial Lidar: Suggestions and Novel Techniques Based on Comparisons Between Scanners and Forest Treatments. Earth and Space Science, 5(11), 753–776. 10.1029/2018EA000417

Dunn, O. J. (1964). Multiple Comparisons Using Rank Sums. Technometrics, 6(3), 241–252. 10.1080/00401706.1964.10490181

Ehbrecht, M., Schall, P., Juchheim, J., Ammer, C., & Seidel, D. (2016). Effective number of layers: A new measure for quantifying three-dimensional stand structure based on sampling with terrestrial LiDAR. Forest Ecology and Management, 380, 212–223. 10.1016/j.foreco.2016.09.003

Feigl, J., Frey, J., Seifert, T., & Koch, B. (2025). Close-Range Remote Sensing of Forest Structure for Biodiversity Assessments: A Systematic Literature Review. Current Forestry Reports, 11, 18. 10.1007/s40725-025-00251-x

Franklin, J. F., & Van Pelt, R. (2004). Spatial aspects of structural complexity in old-growth forests. Journal of Forestry, 102(3), 22–29.

Gao, T., Hedblom, M., Emilsson, T., & Nielsen, A. B. (2014). The role of forest stand structure as biodiversity indicator. Forest Ecology and Management, 330, 82–93. 10.1016/j.foreco.2014.07.007

Gauld, J. H. (1982). Native pinewood soils in the northern section of Abernethy forest. Scottish Geographical Magazine, 98(1), 48–56. 10.1080/00369228208736514

Gullett, P. R., Leslie, C., Mason, R., Ratcliffe, P., Sargent, I., Beck, A., Cameron, T., Cowie, N. R., Hetherington, D., MacDonell, T., Moat, T., Moore, P., Teuten, E., & Hancock, M. H. (2023). Woodland expansion in the presence of deer: 30 years of evidence from the Cairngorms Connect landscape restoration partnership. Journal of Applied Ecology, 60(11), 2298–2308. 10.1111/1365-2664.14501

Holvoet, J., Eichhorn, M. P., Giannetti, F., Kükenbrink, D., Liang, X., Mokroš, M., Novotný, J., Pitkänen, T. P., Puliti, S., Skudnik, M., Stereńczak, K., Terryn, L., Vega, C., & Torresan, C. (2025). Terrestrial and mobile laser scanning for national forest inventories: From theory to implementation. Remote Sensing of Environment, 329, 114947. 10.1016/j.rse.2025.114947

Kent Wildlife Trust. (n.d.). The Wilder Blean Project | Kent Wildlife Trust. Retrieved 15 July 2026, from https://www.kentwildlifetrust.org.uk/projects/wilder-blean

Khosravipour, A., Skidmore, A. K., Isenburg, M., Wang, T., & Hussin, Y. A. (2014). Generating Pit-free Canopy Height Models from Airborne Lidar. Photogrammetric Engineering & Remote Sensing, 80(9), 863–872. 10.14358/PERS.80.9.863

Kirby, K. J. (2004). A model of a natural wooded landscape in Britain as influenced by large herbivore activity. Forestry: An International Journal of Forest Research, 77(5), 405–420. 10.1093/forestry/77.5.405

Kruskal, W. H., & Wallis, W. A. (1952). Use of Ranks in One-Criterion Variance Analysis. Journal of the American Statistical Association, 47(260), 583–621. 10.2307/2280779

Kunz, M. (2019). *tre3d: An R package for computating pinguin metrics* [R Package]. https://rdrr.io/github/spatial-mk/tre3d/src/R/ENL.R

Kuuluvainen, T. (2009). Forest Management and Biodiversity Conservation Based on Natural Ecosystem Dynamics in Northern Europe: The Complexity Challenge. Ambio, 38(6), 309–315. https://www.jstor.org/stable/40390241

Lang, A. K., LaRue, E. A., Kivlin, S. N., Edwards, J. D., Phillips, R. P., Gallion, J., Kong, N., Parker, J. D., McCormick, M. K., Domke, G. M., & Fei, S. (2023). Forest structural diversity is linked to soil microbial diversity. Ecosphere, 14, 14e4702. 10.1002/ecs2.4702

Larsen, J. B., Angelstam, P., Bauhus, J., Carvalho, J. F., Diaci, J., Dobrowolska, D., Gazda, A., Gustafsson, L., Krumm, F., Knoke, T., Konczal, A., Kuuluvainen, T., Mason, B., Motta, R., Pötzelsberger, E., Rigling, A., & Schuck, A. (2022). *Closer-to-Nature Forest Management* [From Science to Policy]. European Forest Institute. 10.36333/fs12

LaRue, E. A., Fahey, R., Fuson, T. L., Foster, J. R., Matthes, J. H., Krause, K., & Hardiman, B. S. (2022). Evaluating the sensitivity of forest structural diversity characterization to LiDAR point density. Ecosphere, 13(9), e4209. 10.1002/ecs2.4209

LaRue, E. A., Fahey, R. T., Alveshere, B. C., Atkins, J. W., Bhatt, P., Buma, B., Chen, A., Cousins, S., Elliott, J. M., Elmore, A. J., Hakkenberg, C. R., Hardiman, B. S., Johnson, J. S., Kashian, D. M., Koirala, A., Papeş, M., St Hilaire, J. B., Surasinghe, T. D., Zambrano, J., … Fei, S. (2023). A theoretical framework for the ecological role of three-dimensional structural diversity. Frontiers in Ecology and the Environment, 21(1), 4–13. 10.1002/fee.2587

LaRue, E. A., Hardiman, B. S., Elliott, J. M., & Fei, S. (2019). Structural diversity as a predictor of ecosystem function. Environmental Research Letters, 14(11), 114011. 10.1088/1748-9326/ab49bb

Li, S., Dai, L., Wang, H., Wang, Y., He, Z., & Lin, S. (2017). Estimating Leaf Area Density of Individual Trees Using the Point Cloud Segmentation of Terrestrial LiDAR Data and a Voxel-Based Model. Remote Sensing, 9(11). 10.3390/rs9111202

Lines, E. R., Fischer, F. J., Owen, H. J. F., & Jucker, T. (2022). The shape of trees: Reimagining forest ecology in three dimensions with remote sensing. Journal of Ecology, 110(8), 1730–1745. 10.1111/1365-2745.13944

Loke, L. H. L., & Chisholm, R. A. (2022). Measuring habitat complexity and spatial heterogeneity in ecology. Ecology Letters, 25(10), 2269–2288. 10.1111/ele.14084

MacArthur, R. H., & MacArthur, J. W. (1961). On Bird Species Diversity. Ecology, 42(3), 594–598. 10.2307/1932254

Martin, M., Shorohova, E., & Fenton, N. J. (2023). Embracing the Complexity and the Richness of Boreal Old-Growth Forests: A Further Step Toward Their Ecosystem Management. In M. M. Girona, H. Morin, S. Gauthier, & Y. Bergeron (Eds), Boreal Forests in the Face of Climate Change: Sustainable Management (pp. 191–218). Springer International Publishing. 10.1007/978-3-031-15988-6_7

Mason, W. L., Connolly, T., Pommerening, A., & Edwards, C. (2007). Spatial structure of semi-natural and plantation stands of Scots pine (Pinus sylvestris L.) in northern Scotland. Forestry: An International Journal of Forest Research, 80(5), 567–586. 10.1093/forestry/cpm038

McElhinny, C., Gibbons, P., Brack, C., & Bauhus, J. (2005). Forest and woodland stand structural complexity: Its definition and measurement. Forest Ecology and Management, 218(1), 1–24. 10.1016/j.foreco.2005.08.034

Melin, M., Hinsley, S. A., Broughton, R. K., Bellamy, P., & Hill, R. A. (2018). Living on the edge: Utilising lidar data to assess the importance of vegetation structure for avian diversity in fragmented woodlands and their edges. Landscape Ecology, 33(6), 895–910. 10.1007/s10980-018-0639-7

Mönkkönen, M., Brazaitis, G., Brūmelis, G., Jonsson, B.-G., Lõhmus, A., Mäkipää, R., & Syrjänen, K. (2026). From guidelines to practice: Operational criteria for identifying old-growth forests in northern Europe. Ambio. 10.1007/s13280-026-02457-0

Morales, A., & MacFarlane, D. W. (2025). *rTwig*: An R package to correct overestimated small branches and twigs in quantitative structure models of trees. Science of Remote Sensing, 12, 100284. 10.1016/j.srs.2025.100284

NatureScot. (2023, March 2). A guide to understanding the Scottish Ancient Woodland Inventory (AWI). https://www.nature.scot/doc/guide-understanding-scottish-ancient-woodland-inventory-awi

Peterken, G. (1983). Peterken: Woodland conservation in Britain. Conservation in Perspective, 83–100. https://scholar.google.com/scholar_lookup?title=Woodland%20conservation%20in%20Britain&publication_year=1983&author=G.F.%20Peterken

Pimont, F., Allard, D., Soma, M., & Dupuy, J.-L. (2018). Estimators and confidence intervals for plant area density at voxel scale with T-LiDAR. Remote Sensing of Environment, 215, 343–370. 10.1016/j.rse.2018.06.024

Pommerening, A. (2002). Approaches to quantifying forest structures. Forestry: An International Journal of Forest Research, 75(3), 305–324. 10.1093/forestry/75.3.305

Pratt, S. (2007). Long term landscape dynamics in a Caledonian pine forest. https://era.ed.ac.uk/handle/1842/7123

R Core Team. (2025). *R: A language and environment for statistical computing.* R Foundation for Statistical Computing. https://www.r-project.org/

Rappa, N. J., Staab, M., Ruppert, L.-S., Frey, J., Bauhus, J., & Klein, A.-M. (2023). Structural elements enhanced by retention forestry promote forest and non-forest specialist bees and wasps. Forest Ecology and Management, 529, 120709. 10.1016/j.foreco.2022.120709

Regnery, B., Couvet, D., Kubarek, L., Julien, J.-F., & Kerbiriou, C. (2013). Tree microhabitats as indicators of bird and bat communities in Mediterranean forests. Ecological Indicators, 34, 221–230. 10.1016/j.ecolind.2013.05.003

Rosen, A., Jörg Fischer, F., Coomes, D. A., Jackson, T. D., Asner, G. P., & Jucker, T. (2024). Tracking shifts in forest structural complexity through space and time in human-modified tropical landscapes. Ecography, 2024(11), e07377. 10.1111/ecog.07377

Ross, C. W., Loudermilk, E. L., O’Brien, J. J., & Snitker, G. (2024). Lidar-derived structural-complexity data across four experimental forests. Data in Brief, 57, 110955. 10.1016/j.dib.2024.110955

Roussel, J.-R., Auty, D., Coops, N. C., Tompalski, P., Goodbody, T. R. H., Meador, A. S., Bourdon, J.-F., de Boissieu, F., & Achim, A. (2020). lidR: An R package for analysis of Airborne Laser Scanning (ALS) data. Remote Sensing of Environment, 251, 112061. 10.1016/j.rse.2020.112061

Saarinen, N., Calders, K., Kankare, V., Yrttimaa, T., Junttila, S., Luoma, V., Huuskonen, S., Hynynen, J., & Verbeeck, H. (2021). Understanding 3D structural complexity of individual Scots pine trees with different management history. Ecology and Evolution, 11(6), 2561–2572. 10.1002/ece3.7216

Schooler, S. L., & Zald, H. S. J. (2019). Lidar Prediction of Small Mammal Diversity in Wisconsin, USA. Remote Sensing, 11(19), 2222. 10.3390/rs11192222

Scottish Government. (2026). Scottish Land LiDAR Programme—2025 capture—LAS - data.gov.uk. https://ckan.publishing.service.gov.uk/dataset/scottish-land-lidar-programme-2025-capture-las

Seidel, D. (2018). A holistic approach to determine tree structural complexity based on laser scanning data and fractal analysis. Ecology and Evolution, 8(1), 128–134. 10.1002/ece3.3661

Silva, C. A., Valbuena, R., Pinagé, E. R., Mohan, M., de Almeida, D. R. A., North Broadbent, E., Jaafar, W. S. W. M., de Almeida Papa, D., Cardil, A., & Klauberg, C. (2019). ForestGapR: An r Package for forest gap analysis from canopy height models. Methods in Ecology and Evolution, 10(8), 1347–1356. 10.1111/2041-210X.13211

Steven, H. M., & Carlisle, A. (1959). The Native Pinewoods of Scotland. Oliver and Boyd.

Stiers, M., Annighöfer, P., Seidel, D., Willim, K., Neudam, L., & Ammer, C. (2020). Quantifying the target state of forest stands managed with the continuous cover approach – revisiting Möller’s “Dauerwald” concept after 100 years. Trees, Forests and People, *1*, 100004. 10.1016/j.tfp.2020.100004

Summers, R. (2018). Abernethy Forest: The History and Ecology of an Old Scottish Pinewood. Centre of Conservation Science, RSPB. https://www.nhbs.com/abernethy-forest-book

Summers, R. W., Proctor, R., Raistrick, P., & Taylor, S. (1997). The structure of Abernethy Forest, Strathspey, Scotland. Botanical Journal of Scotland, 49(1), 39–55. 10.1080/03746609708684851

Van Ewijk, K. Y., Treitz, P. M., & Scott, N. A. (2011). Characterizing Forest Succession in Central Ontario using Lidar-derived Indices. Photogrammetric Engineering & Remote Sensing, 77(3), 261–269. 10.14358/PERS.77.3.261

Vera, F. W. M., Bakker, E. S., & Olff, H. (2006). Large herbivores: Missing partners of western European light-demanding tree and shrub species? In K. Danell, R. Bergström, P. Duncan, & J. Pastor (Eds), Large Herbivore Ecology, Ecosystem Dynamics and Conservation (1st edn, pp. 203–231). Cambridge University Press. 10.1017/CBO9780511617461.009

Wildermuth, B., Dönges, C., Matevski, D., Penanhoat, A., Seifert, C. L., Seidel, D., Scheu, S., & Schuldt, A. (2023). Tree species identity, canopy structure and prey availability differentially affect canopy spider diversity and trophic composition. Oecologia, 203(1), 37–51. 10.1007/s00442-023-05447-1

Wildermuth, B., Penanhoat, A., Sennhenn-Reulen, H., Matevski, D., Drescher, J., Aubry-Kientz, M., Seidel, D., & Schuldt, A. (2024). Canopy structure influences arthropod communities within and beyond tree identity effects: Insights from combining LiDAR data, insecticidal fogging and machine learning regression modelling. Ecological Indicators, 160, 111901. 10.1016/j.ecolind.2024.111901

Wilkes, P. (2023). *Philwilkes/rxp-pipeline* [Python]. https://github.com/philwilkes/rxp-pipeline (Original work published 2021)

Wilkes, P., Lau, A., Disney, M., Calders, K., Burt, A., Gonzalez de Tanago, J., Bartholomeus, H., Brede, B., & Herold, M. (2017). Data acquisition considerations for Terrestrial Laser Scanning of forest plots. Remote Sensing of Environment, 196, 140–153. 10.1016/j.rse.2017.04.030

Willim, K., Stiers, M., Annighöfer, P., Ehbrecht, M., Ammer, C., & Seidel, D. (2020). Spatial Patterns of Structural Complexity in Differently Managed and Unmanaged Beech-Dominated Forests in Central Europe. Remote Sensing, 12, 1907. 10.3390/rs12121907

Zeller, L., Förster, A., Keye, C., Meyer, P., Roschak, C., & Ammer, C. (2023). What does literature tell us about the relationship between forest structural attributes and species richness in temperate forests? – A review. Ecological Indicators, 153, 110383. 10.1016/j.ecolind.2023.110383

Zellweger, F., Braunisch, V., Baltensweiler, A., & Bollmann, K. (2013). Remotely sensed forest structural complexity predicts multi species occurrence at the landscape scale. Forest Ecology and Management, 307, 303–312. 10.1016/j.foreco.2013.07.023

Zhang, W., Qi, J., Wan, P., Wang, H., Xie, D., Wang, X., & Yan, G. (2016). An Easy-to-Use Airborne LiDAR Data Filtering Method Based on Cloth Simulation. Remote Sensing, 8(6), 501. 10.3390/rs8060501

Zhang, Z., Cao, L., & She, G. (2017). Estimating Forest Structural Parameters Using Canopy Metrics Derived from Airborne LiDAR Data in Subtropical Forests. Remote Sensing, 9(940), Article 9. 10.3390/rs9090940

