## Supporting Information for "Simplifying complexity: the reliability of lidar-derived forest structural complexity metrics for conservation management"

**Table S1.** Interpretation of the forest structural complexity metrics used in this study, and the relationships reported for them in the literature. Metric groups are adapted from those described by Feigl et al. (2025).

| Metric Group | Metric name | Interpretation | A study relating the metric to forest management or biodiversity |
|---|---|---|---|
| Density | (1) Gap Fraction (3D) | Proportion of empty voxels; inverse of space filling, indicating light penetration | La Cava et al. 2024; Zellweger et al. 2013; Wildermuth et al. 2024 |
| | (2) Vertical space filling | Higher values reflect canopy packing and resource use efficiency | Georgi et al. 2022; Ray et al. 2024; Wildermuth et al. 2024; Shokirov et al. 2024; Juchheim et al. 2017; |
| Density variability | Gini coefficient (space filling) (3, 4) | Inequality in spatial distribution of vegetation; high values indicate clustering (vertical and horizontal) | Stiers et al. 2020 |
| | Coefficient of variation (space filling) (5,6) | Variability in vegetation distribution across vertical or horizontal space | Stiers et al. 2020 |
| Density layering | CV of LAD (7) | Variability in leaf area density across height; higher values indicates vertical heterogeneity | Carrasco et al. 2019; Bouvier et al. 2017; |
| | ENL (Effective Number of Layers) (8) | Number of distinct vertical layers accounting for their relative importance. A higher ENL, the more effective layers | Wildermuth et al. 2024; Wildermuth et al. 2023; Rappa et al. 2022; Rappa et al. 2023; Ehbrecht et al. 2016 |
| | FHD Foliage Height Diversity (9) | Evenness of vegetation across vertical layers; higher values indicate more evenly distributed canopy | MacArthur & MacArthur 1961; Somveille et al. 2025 |
| | Vertical Complexity Index (VCI) (10) | Shannon entropy across height bins, normalised by maximum entropy; evenness of vertical distribution. Vertical only. | Shokirov et al. 2024; Adiningrat et al. 2025 |
| Fractal | Box dimension (BoxD) (11) | Fractal measure of structural complexity; reflects how completely vegetation fills space | Wildermuth et al. 2024; Wildermuth et al. 2023; Stiers et al. 2018; Neudam et al. 2022; Cimdins et al. 2025 |
| Height | Max (height) (12) | Maximum canopy height; reflects structural development | Coverdale et al. 2024; Bouvier et al. 2017 |
| | Mean (height) (13) | Average canopy height; indicates overall stand stature | Blakey et al. 2017; St. Rose & Naithani 2025; Davison et al. 2023 |
| | SD (height) (14) | Variability in canopy height; higher values reflects more vertical heterogeneity | Blakey et al. 2017; Hendel et al. 2025; Davison et al. 2023; Carrasco et al. 2019 |
| | CV of canopy height *(CV_height) (15)* | Standard deviation of canopy height divided by the mean; standardised for stature, so comparable across stands. | Coverdale et al. 2024; Wildermuth et al. 2024; Shokirov et al. 2024; |

| | | | |
|---|---|---|---|
| | Gini coefficient of canopy height *(Gini_height)* (16) | Inequality of canopy height across canopy height model pixels; a few tall pixels among many low ones. | Bouvier et al. 2017 |
| Rugosity | Rumple index (17) | Ratio of canopy surface area to ground area; indicates canopy roughness | St. Rose & Naithani 2025; Coverdale et al. 2024; Wildermuth et al. 2024; Adiningrat et al. 2025 |
| | Rugosity (18) | Variation in canopy surface height; higher values reflect greater structural irregularity | St. Rose & Naithani 2025; Coverdale et al. 2024 |
| Openness | Gap fraction (2D) (19) | Higher values indicate a larger number of gaps, reflecting light availability | St. Rose & Naithani 2025; Wildermuth et al. 2024; Rappa et al. 2023; Adiningrat et al. 2025 |
| | Canopy Cover (20) | Proportion of ground area covered by canopy; higher values indicate canopy closure but not necessarily greater complexity, as dense monocultures can achieve high canopy cover | Bradley et al. 2022; Coverdale et al. 2024; St. Rose & Naithani 2025; Blakey et al. 2017; |
| | Gap area CV (21) | Higher values represent higher variability of gap size and therefore a heterogenous distribution of light | Wildermuth et al. 2024 |

Notes
Cimdins et al. (2025) compute the Gini coefficient on segmented individual-tree heights rather than canopy height model pixels, gap fraction as the proportion of returns below 2 m rather than below a relative canopy-height threshold, and box dimension per tree rather than per stand;
Space filling appears in the literature under several names — vegetation volume, canopy volume, voxel occupancy, vegetation area index, canopy packing, canopy space occupation, and the filled and empty voxel classes of Lefsky et al. (1999). Entries were matched across these terms.
Dash indicates that no study was found.
Numbers relate to in text referencing of metrics

**Table S3.** Examples of hyperparameter use in the literature.

| Metric | Author | | Hyper-parameter | Hyperparameter size (m) |
|---|---|---|---|---|
| Cover | Li 2022 | TLS | chm | 0.01 |
| | Muir et al., 2018 | TLS | raster | 0.1 |
| | Coverdale 2024 | UAV | chm | 0.25 |
| | Wildermuth 2024 | MLS | chm | 0.5 |
| | Blakey et al 2017 | TLS | raster | 0.5 |
| | Penman 2023 | TLS | raster | 0.2 |
| Space filling/Voxel Occupancy/PAI | Neudam 2022 | MLS + TLS | voxel | 0.2 |
| | Aalto 2023 | TLS | voxel | 1 |
| | Muir et al., 2018 | TLS | voxel | 0.1 |
| | Flynn et al 2023 | TLS | voxel | 0.05 |
| FHD | Somveille 2025 | MLS | voxel | 1 |
| | Aalto 2023 | TLS | voxel | 1 |
| | Penman 2023 | TLS | voxel | 0.1 |
| Canopy height | Blakey et al 2017 | TLS | raster | 0.5 |
| | Li 2022 | TLS | chm | 0.01 |
| | Shokirov 2023 | TLS + UAV | point cloud | 0.01 |
| | Coverdale 2024 | UAV | raster | 0.25 |
| | Wildermuth 2024 | MLS | voxel | 0.5 |
| | Shokirov 2024 | TLS | point cloud | 0.01 |
| | Penman 2023 | TLS | raster | 0.2 |
| Standard deviation of height | Shokirov 2023 | TLS + UAV | point cloud | 0.01 |
| | Blakey et al 2017 | TLS | raster | 0.05 |
| Vegetation density | Willim 2020 | TLS | voxel | 0.2 |
| | Shokirov 2023 | TLS + UAV | grid | 0.25 |
| | Eichhorn 2017 | TLS | voxel | 0.01 |
| | Penman 2023 | TLS | voxel | 0.1 |
| Gap volume | Sara La Cava 2024 | TLS | voxel | 0.25 |
| | Atkins 2018 | MLS | voxel | 1 |
| | Blakey et al 2017 | TLS | voxel | 0.25 x 5 x 5 |
| | Penman 2023 | TLS | voxel | 0.1 |
| Rugosity | Coverdale 2024 | UAV | chm | 0.25 |
| | Blakey et al 2017 | TLS | raster | 0.5 |
| | Blakey et al 2017 | TLS | raster | 0.5 |
| Correlation Variation of height | Coverdale 2024 | UAV | chm | 0.25 |
| | Shokirov 2024 | TLS | point cloud | 0.01 |
| | Wildermuth 2024 | MLS | chm | 0.5 |
| Vegetation volume | Shokirov 2024 | TLS | voxel | 0.5 |
| | Wildermuth 2024 | MLS | voxel | 0.5 |
| | Shokirov 2023 | TLS + UAV | voxel | 0.5 |
| VAI/LAI | Li 2022 | TLS | Height bin | 0.5 |
| | Atkins 2018 | MLS | voxel | 1 |

| | | | | |
|---|---|---|---|---|
| | Somveille 2025 | MLS | voxel | 1 |
| Leaf area density | Somveille 2025 | MLS | voxel | 1 |
| | Blakey et al 2017 | TLS | pillar | 0.25 x 5 x 5 |
| Crown volume | Korhonen et al. 2011 | ALS | dsm | 0.5 |
| | Brede et al. (2017) | TLS | chm | 0.5 |
| | Liang (2019) | TLS | voxel | 0.5 |
| SSCI | Ebrecht *et al.,* 2017 | TLS | voxel + height bin | 0.2 in 1m layers |
| | Greve 2018 | TLS | voxel + height bin | 0.2 voxels, 1m height bin (literary standard Ebrecht et al., 2017) |
| | Willim 2022 | TLS | voxel + height bin | 0.2 voxels, 1m height bin (literary standard Ebrecht et al., 2017) |
| | Anika Goßmann 2023 | TLS | voxel + height bin | 0.2 voxels, 1m height bin (literary standard Ebrecht et al., 2017) |
| Rumple Index | Wildermuth 2024 | MLS | chm | 0.5 |
| | Penman 2023 | TLS | raster | 0.2 |
| | Coverdale 2024 | UAV | chm | 0.25 |
| Box dimension | Willim 2020 | TLS | point cloud | 0.2 |
| | Wildermuth 2024 | MLS | point cloud | 0.05 |
| | Neudam 2022 | MLS | point cloud | 0.2 |
| Rumple Index Understory | Penman 2023 | TLS | raster | 0.01 |
| Skewness of height | Shokirov 2023 | TLS + UAV | point cloud | 0.01 |
| | Shokirov 2024 | TLS | point cloud | 0.01 |
| Kurtosis of height | Shokirov 2023 | TLS + UAV | point cloud | 0.01 |
| | Pickering 2025 | GEDI | | |
| | Shokirov 2024 | TLS | point cloud | 0.01 |
| VCI | Adiningrat 2025 | | | |
| | Shokirov 2024 | TLS | point cloud | 0.5 |
| Gap Fraction | Shokirov 2024 | TLS | point cloud | 0.01 |
| Canopy roughness | Shokirov 2024 | TLS | point cloud | 0.01 |
| Canopy Shannon | Shokirov 2024 | TLS | point cloud | 0.01 |
| Mean Gap Area | Wildermuth 2024 | MLS | chm | 0.5 |
| Coefficient of variation Gap Area | Wildermuth 2024 | MLS | chm | 0.5 |
| Coefficient of variation of intra-canopy gap height | Wildermuth 2024 | MLS | voxel | 0.5 |
| Coefficient of variation of intra-canopy gap volume | Wildermuth 2024 | MLS | voxel | 0.5 |
| ENL | Wildermuth 2024 | MLS | raster | 0.5 |
| | Willim 2022 | TLS | voxel + height bin | 0.2 voxels, 1m height bin (literary standard Ebrecht et al., 2017) |
| PAD | Aalto 2023 | TLS | voxel | 1 |

**Table S4.** Results of maximum mean separation analysis to assist hyperparameter choice

| metric | group | resolution | separation |
|---|---|---|---|
| CVxy | Density | 0.02 | 3.876786671 |
| CVz | Density | 0.02 | 9.395494166 |
| gap_frac_2D | Openness | 0.02 | 1.964262291 |
| rumple | Rugosity | 0.02 | 47.97246034 |
| Gini_height | Rugosity | 0.02 | 1.546055157 |
| CV_height | Rugosity | 0.05 | 452.3576339 |
| Canopy cover | Openness | 0.05 | 2.229959397 |
| sd_height | Rugosity | 0.1 | 14.38913883 |
| SFz | Density | 0.2 | 44.17423308 |
| max_height | Height | 0.2 | 47.39874524 |
| rugosity | Rugosity | 0.5 | 3.55407138 |
| CV_LAD | Density | 0.7 | 1.177934343 |
| GINIxy | Density | 0.75 | 0.463627237 |
| CV_gap_area | Openness | 1 | 5.718422134 |
| ENL | Density | 1.02 | 40.5463041 |
| FHD | Density | 1.35 | 4.397076513 |
| GINIz | Density | 1.45 | 0.949160207 |
| VCI | Density | 1.45 | 1.202439004 |
| gap_frac_3D | Density | 1.45 | 1.041267319 |
| mean_height | Height | 1.45 | 47.73772659 |

a)

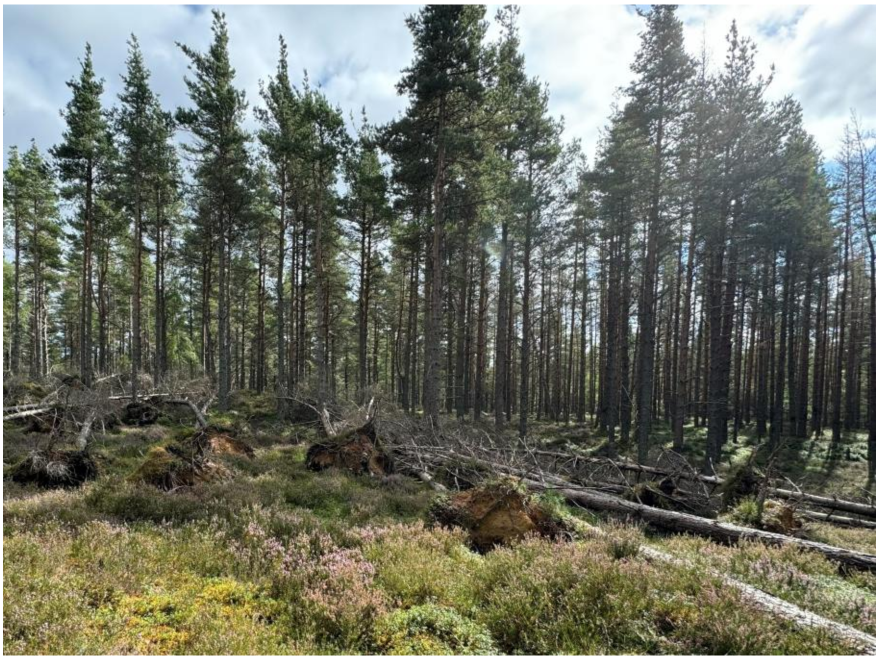

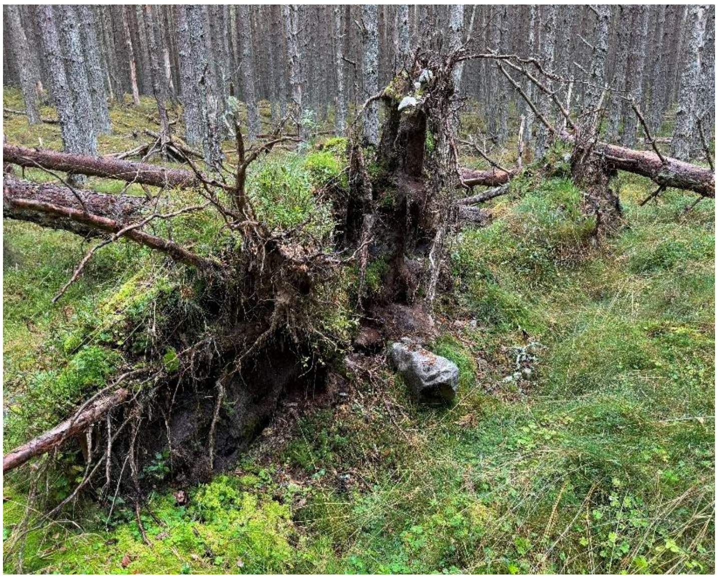

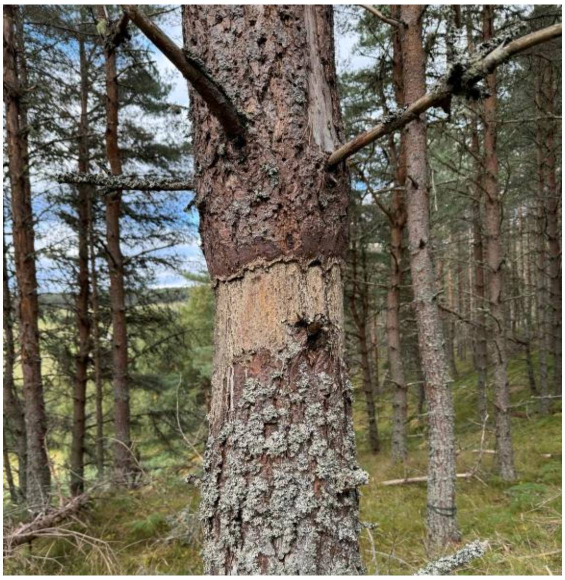

b)

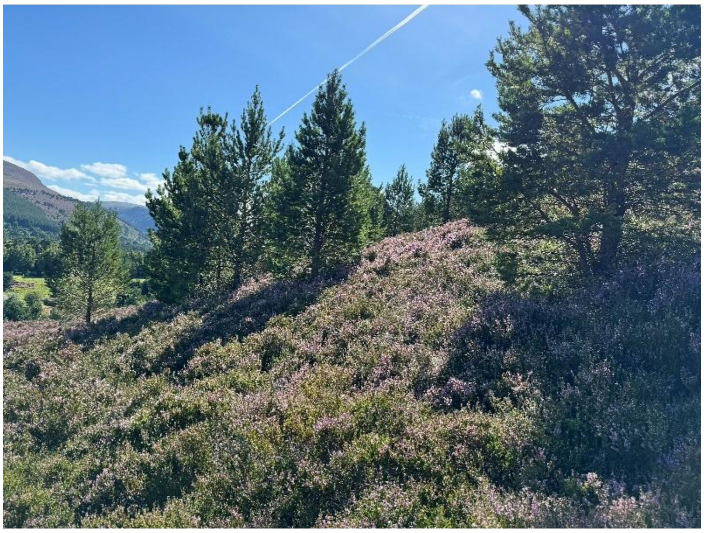

c)

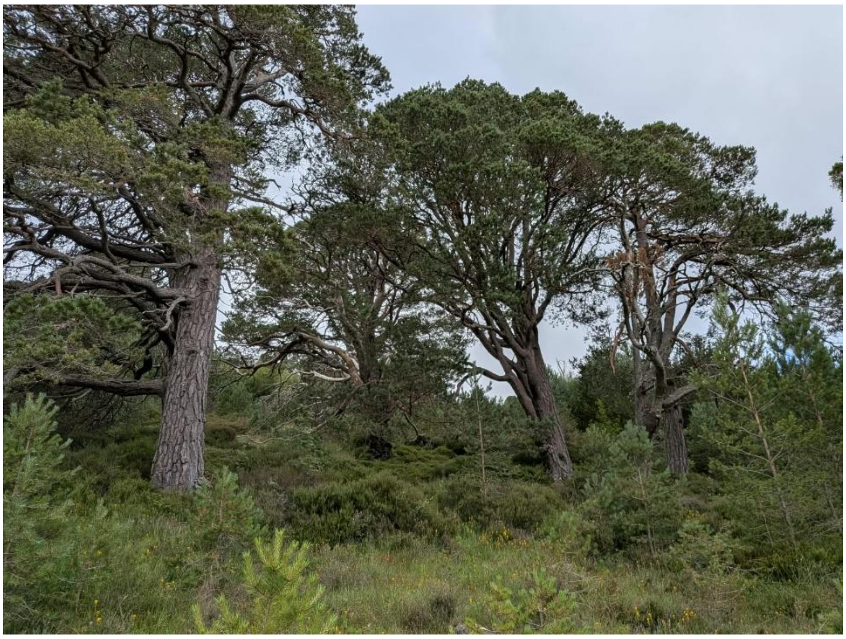

d)

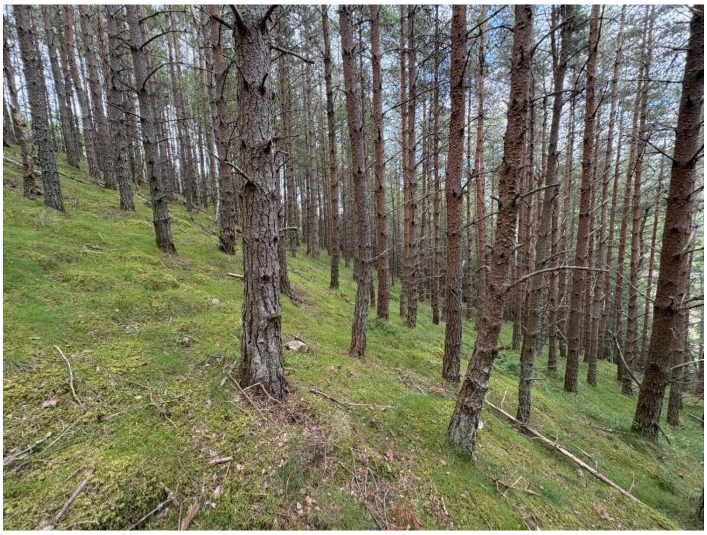

Figure S1 Photos showing examples of the different structural categories. a) Old plantation managed for conservation via re-structuring, b) natural regeneration, c) Ancient woodland and d) Old plantation
